# CDDO-Imidazole regulates RBC alloimmunization to the KEL antigen by activating Nrf2

**DOI:** 10.1101/2025.04.03.645598

**Authors:** Che-Yu Chang, Rosario Hernández-Armengol, Kausik Paul, June Young Lee, Karina Nance, Tomohiro Shibata, Peibin Yue, Christian Stehlik, David R. Gibb

## Abstract

During red blood cell (RBC) transfusion, production of alloantibodies can promote significant hemolytic events. However, most transfusion recipients do not form anti-RBC alloantibodies. Identifying mechanisms that inhibit alloimmunization may lead to prophylactic interventions. One potential regulatory mechanism is activation of the transcription factor, nuclear factor erythroid-derived 2-like 2 (Nrf2), a master regulatory of antioxidant pathways. Pharmacologic Nrf2 activators improve sequelae of sickle cell disease in pre-clinical models. The Nrf2 activator, 1-[2-cyano-3-,12-dioxooleana-1,9(11)-dien-28-oyl]imidazole (CDDO-Im), suppresses production of inflammatory cytokines including type 1 interferons (IFNα/β), which have been implicated in promoting RBC alloimmunization in transfusion models. Thus, we tested the hypothesis that the Nrf2 activator, CDDO-Im, regulates RBC alloimmunization. Here, we report that CDDO-Im induced Nrf2 activated gene expression and suppressed poly(I:C)-induced IFNα/β-stimulated gene (ISG) expression in human macrophages and murine blood leukocytes. In addition, following transfusion of wildtype mice with RBCs expressing the KEL antigen, CDDO-Im treatment inhibited poly(I:C)-induced anti-KEL IgG production and promoted post-transfusion recovery of KEL+ RBCs, but failed to do so in *Nrf2*^-/-^ mice. Results indicate that activation of the Nrf2 antioxidant pathway regulates RBC alloimmunization to the KEL antigen in a pre-clinical model. If findings translate to other models and human studies, Nrf2 activators may represent a potential prophylactic intervention to inhibit alloimmunization.

**Key Points:**

- The antioxidant pathway, Nrf2, inhibits anti-RBC alloantibody responses in a pre-clinical transfusion model.
- Nrf2 activation may represent a prophylactic strategy to inhibit RBC alloimmunization in transfusion recipients.

## Introduction

During red blood cell (RBC) transfusion, most non-ABO antigens are not routinely matched between donors and recipients. This exposure of mis-matched antigens can cause the production of anti-RBC alloantibodies which mediate hemolytic events including hemolytic transfusion reactions, which are a cause of transfusion-associated mortality^1,2^. Transfusion-dependent patients commonly produce alloantibodies against multiple RBC antigens. Acquiring compatible RBCs lacking many RBC antigens for these patients can be difficult if not unachievable. Thus, they commonly experience anemia-induced morbidities and/or receive incompatible RBC transfusions in the presence of anti-RBC antibodies, which can cause hemolytic transfusion reactions^3,4^.

Avoiding RBC antigen exposure is the only clinically used strategy to prevent RBC alloimmunization in transfusion recipients. The provision of extended antigen matched RBCs (i.e. matching of C, E, and KEL antigens in addition to ABO/Rh(D)) has significantly reduced the frequency of alloimmunization in patients with hemoglobinopathies^5^. However, extended antigen matching is not utilized universally^6^. In addition, as there are as many as 340 antigens on the RBC surface and there are numerous variants of the Rh antigens (D, C, E)^7,8^, other strategies to inhibit alloimmunization are needed.

One factor that influences RBC alloimmunization is inflammation in the transfusion recipient. Patients with disseminated viral infections or inflammatory autoimmune diseases have elevated frequencies of alloimmunization^9-13^. In addition, patients with SCD transfused during a vaso occlusive crisis or acute chest syndrome have profoundly increased odds of alloimmunization, compared to patients with SCD in their baseline state of health^5^. We and others have used pre-clinical murine models to investigate mechanisms of inflammation-induced alloimmunization. Viral infection and treatment with the viral mimetic, polyinosinic:polycytidylic acid (poly(I:C)), before transfusion enhances RBC alloimmunization in murine transfusion models^14-17^. We have reported that viral-induced type 1 interferons (IFNα/β) are required for alloimmunization in multiple murine transfusion models^16-19^, and recombinant IFNα (rIFNα) is sufficient to induce alloimmunization^17^.

Unlike patients with inflammation, the incidence of alloimmunization in the general transfused population is low. RBC alloimmunization only occurs in 3-10% of patients in US hospitals^10^. This extends to transfusion-dependent patients with higher frequencies of alloimmunization. While 30-50% of patients with SCD produce RBC alloantibodies, many recipients with SCD do not, despite having a high transfusion burden^10,20^. This indicates that there may be negative regulatory mechanisms that prevent alloimmunization. Such mechanisms could be leveraged for prophylactic interventions to prevent alloimmunization.

One potential regulatory mechanism is activation of the transcription factor, nuclear factor erythroid-derived 2-like 2 (Nrf2), which is a master regulator of antioxidant pathways activated by oxidative stress. While Nrf2 is recognized for its regulatory role in tumor cell proliferation and invasion^21^, its role in transfusion is not known. At steady state, Nrf2 is associated with its principal negative regulator, Kelch-like ECH-associated protein 1 (Keap1), in the cytoplasm. During oxidative stress, Nrf2 dissociates from Keap1, translocates to the nucleus, and induces expression of antioxidant enzymes, including heme oxygenase 1 (HMOX1) and NAD(P)H quinone dehydrogenase 1 (NQO1). Nrf2 also regulates iron metabolism and inflammatory responses^22^. Nrf2 has been shown to suppress inflammatory responses in murine models of inflammation, including sepsis^23^, neuroinflammation^24^, hepatic disease^25^ and viral infection^26^, and there is growing evidence that Nrf2 inhibits IFNα/β responses. Gunderstofte et al. reported that Nrf2 down-regulation or deficiency in bone marrow-derived macrophages (BMDMs) results in elevated IFNα/β and expression of IFNα/β-stimulated gene (ISGs) that are protective in a model of herpes simplex virus-2^26^. A similar inverse correlation between Nrf2 activation and ISG expression was observed in human epithelial cells treated with Nrf2 siRNA^27^.

Due to the antioxidant and anti-inflammatory roles of Nrf2, pharmacologic Nrf2 activators have been tested in many inflammatory conditions. Some have entered clinical trials and are FDA approved for specific indications^28,29^. Most activators are electrophilic compounds that alter cysteines of Keap1, allowing release and activation of Nrf2^30^. CDDO-Im (1-[2-cyano-3-,12-dioxooleana-1,9(11)-dien-28-oyl] imidazole) is a synthetic triterpenoid derived from oleanolic acid and a potent Nrf2 activator^31^. Nrf2 activators, including CDDO-Im, sulforaphane, and dimethyl fumarate (DMF), were shown to inhibit vaso occlusion and vascular inflammation in models of SCD^32-34^. However, a role for Nrf2 activators in regulating RBC alloimmunization has not been previously investigated. Here we investigated the role of CDDO-Im induced Nrf2 activation in regulating RBC alloimmunization in a pre-clinical transfusion model.

## Methods

### CDDO-Im administration to mice

C57BL/6J and *Nrf2*^-/-^ mice were purchased from the Jackson Laboratories (Bar Harbor, ME, USA). K1 transgenic mice expressing the human KEL glycoprotein containing the KEL1 antigen on RBCs were previously described^17^. C57BL/6J and *Nrf2*^-/-^ mice, 8-12 weeks of age, were intraperitoneally injected with 2.5 to 10 mg/kg of CDDO-Im (Tocris Bioscience, Bristol, UK). All animal procedures were approved by the Cedars-Sinai Institutional Animal Care and Use Committee.

### RBC transfusion

K1 and wildtype (WT) RBCs were collected from K1 and C57BL/6 mice, respectively, by phlebotomy and anticoagulated with 12% Citrate Phosphate Dextrose Adenine (CPDA-1, Jorgensen Labs, Melville, NY, USA). RBCs were leukoreduced with leukoreduction syringe filters (Pall, East Hills, NY, USA) and resuspended in PBS. 50 µL of packed RBCs, the approximate murine equivalent of one unit of human RBCs, were transfused via retroorbital or tail-vein injection. Recipient mice were pre-treated with or without CDDO-Im and/or 100 µg polyinosinic: polycytidylic acid (poly(I:C), Invivogen, San Diego, CA, USA) by i.p. injection 3-6 hours prior to transfusion.

### Anti-KEL antibody measurement

Serum was collected from transfused mice 5-14 days after transfusion. Anti-KEL IgM and anti-KEL IgG were measured by flow cytometric crossmatch, as previously described^19^, 5 and 7-14 days after transfusion with K1 RBCs, respectively. Briefly, K1 RBCs from non-transfused mice were incubated with transfusion recipient serum, washed, and stained with secondary antibodies, goat anti-mouse IgG APC or IgM FITC (Jackson ImmunoResearch, West Grove, PA). The mean fluorescence intensity (MFI) was measured on a Cytek Northern Lights flow cytometer (Fremont, CA, USA). The adjusted MFI was calculated by subtracting the reactivity of serum with WT RBCs from serum reactivity with K1 RBCs.

### Post-transfusion recovery

Mice previously transfused with K1 RBCs were re-transfused retro-orbitally with a 1:1 mixture of K1 RBCs and WT RBCs 35 days after the initial transfusion. K1 RBCs were labeled with fluorescent 1,1’-dioctadecyl-3,3,3’,3’-tetramethylindocarbocyanine perchlorate (DiI) and WT RBCs were labeled with 3,3’-dioctadecyloxacarbocyanine perchlorate (DiO, Life Technologies, Camarill, CA). DiI and DiO-labeled RBCs remaining in circulation were quantified by flow cytometry 10 min and 1-3 days after transfusion. The ratio of K1:WT RBCs on days 1-3 was plotted as a percentage of the ratio measured 10 min after transfusion as previously described^19^.

### CDDO-Im treatment of human macrophages

Leukoreduction System cones were obtained following apheresis platelet donation by de-identified platelet donors in the Cedars-Sinai Blood Donation Center. Peripheral blood mononuclear cells were isolated by a Ficoll-Paque Premium density gradient (Cytiva, Marlborough, MA, USA) and monocytes were enriched by magnetic negative selection using the EasySep human monocyte isolation kit (StemCell Technologies, Vancouver, CA). Monocytes were differentiated into macrophages with GM-CSF (50 ng/mL) in serum free Macrophage media SFM (Thermo Fisher Scientific, Waltham, MA, USA) containing penicillin-streptomycin (P/S, 10 U/mL) for 5 days. Macrophages were then treated with CDDO-Im (200-800 nM) or sulforaphane (5-10 µM) for 18hrs in complete RPMI containing 10% FBS, 1% L-glutatmine, 1% P/S, 1% NEAA, 1% Sodium Pyruvate, and 1% HEPES (Thermo Fisher Scientific). For some experiments, macrophages were then washed and cultured with 1 µg/mL poly(I:C) in complete RPMI for 3-24 hrs.

### Flow cytometry of human macrophages

Cultured human macrophages were collected using trypsin-EDTA (0.25%, Thermo Fisher). Fc receptors were blocked using human TruStain FcX and then labeled with mouse anti-human Siglec-1 PE, CD14 PerCP, CD64 BV785, CD38 APC and Zombie NIR (Biolegend, San Diego, CA). Cells were then fixed and permeabilized using the Cyto-Fast Fix/Perm Buffer set (Biolegend) and stained with rat anti-mouse HMOX1 (MA1-112), which was conjugated to Alexa Fluor 405 using the Zenon mouse IgG1 labeling kit according to manufacturer’s instructions (Thermo Fisher Scientific). Macrophages were analyzed with a Cytek Northern Lights flow cytometer.

### Quantitative PCR

RNA was isolated from human macrophages and murine blood leukocytes using the Qiagen RNeasy mini-kit (Hilden, DE) and reverse transcribed into cDNA with the Maxima H Minus cDNA Synthesis Master mix (Thermo Fisher Scientific). cDNA encoding mouse *HMOX1, NQO1*, and *GAPDH*, and human *NFE2L2* (Nrf2), *AKR1C1, HMOX1, NQO1, MXA, CXCL10* (IP-10), *ISG15, IFIT3, IRF5, IRF7*, and *GAPDH* was measured using PowerUp SYBR Green master mix on a QuantStudio 5 Real-Time PCR System (Thermo Fisher Scientific). Primer sequences are listed in **Supplemental Table 1**. The relative expression of target genes, compared to *GAPDH*, was determined using Thermo Fisher Connect software.

### Cytokine quantification

Mouse serum cytokines were measured using the LEGENDplex mouse anti-virus response bead assay according to manufacturer’s instructions (Biolegend). Samples were analyzed on the Cytek flow cytometer and calculated using the LEGENDplex Data Analysis Software Suite.

### Statistical Analysis

GraphPad Prism was used for statistical analysis. For anti-RBC antibody and post-transfusion recovery data, non-parametric testing with a Mann-Whitney U test or a Kruskal-Wallis with a Dunn’s post-test was completed for comparing 2 or more than 2 samples, respectively. For qPCR and cytokine data, parametric testing with a Student’s t-test or a one-way ANOVA with a Tukey’s post-test were used for comparing 2 or more than 2 samples, respectively. P-values <0.05 were considered statistically significant. Graphs show the mean as a vertical bar and individual mice or human samples as a white circle.

## Results

### CDDO-Im induces expression of Nrf2-stimulated genes in murine blood leukocytes

CDDO-Im has been shown to be a potent activator of Nrf2 signaling^31^. Thus, we initially verified that CDDO-Im treatment of wildtype (WT) mice, 6 hrs prior to analysis, induces expression of Nrf2-stimulated antioxidant genes, HMOX1 and NQO1, in blood leukocytes. Given that poly(I:C) was utilized in transfusion experiments below, WT mice were also treated with or without poly(I:C) 3 hrs prior to analysis. While poly(I:C) did not alter expression of the Nrf2-inducible genes in peripheral blood cells, addition of CDDO-Im significantly increased expression of HMOX1 and NQO1 (**Supplemental Figure 1**).

### CDDO-Im inhibits RBC alloimmunization in a murine transfusion model

To determine the role of CDDO-Im-mediated Nrf2 activation in RBC alloimmunization, we measured anti-RBC antibody responses in a mouse transfusion model. RBCs collected from transgenic mice expressing the human KEL glycoprotein specifically on RBCs were leukoreduced and transfused into recipient WT mice treated with or without CDDO-Im and/or poly(I:C) 6 hrs and 3 hrs prior to transfusion, respectively (**Figure 1A**). Compared to mice treated with only poly(I:C), mice treated with poly(I:C) and 10 mg/kg CDDO-Im produced lower levels of anti-KEL IgM 5 days after transfusion (**Figure 1B**). In addition, poly(I:C) induced anti-KEL IgG antibody production 7 and 14 days after transfusion, while increasing doses of CDDO-Im significantly inhibited poly(I:C)-induced alloimmunization. Treatment with CDDO-Im, in the absence of poly(I:C), had no effect on antibody responses (**Figure 1C**).

**Figure 1.**
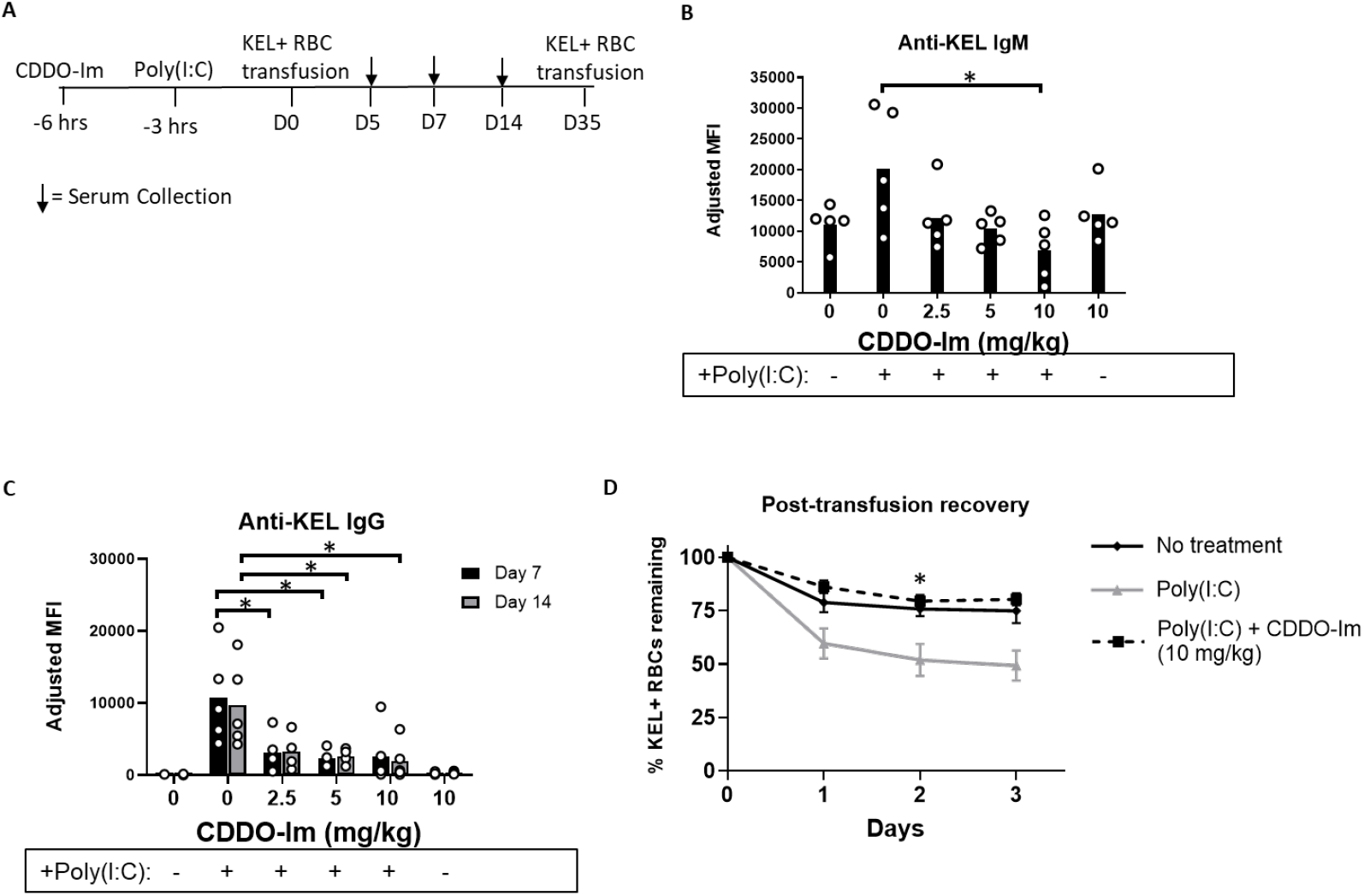
CDDO-Im inhibits inflammation-induced RBC alloimmunization. WT mice were treated with or without poly(I:C) and/or CDDO-Im 3 and 6 hrs before transfusion, respectively, with KEL+ RBCs. (**A**) Timeline of treatments, transfusions, and serum collections. (**B**) Serum anti-KEL IgM levels collected 5 days after transfusion. (**C**) Serum anti-KEL IgG levels collected 7 and 14 days after transfusion. (**D**) Mice were re-transfused with fluorescently labeled KEL+ RBCs and control WT RBCs 35 days after the initial transfusion. Post-transfusion recovery of KEL+ RBCs: WT RBCs ratios 1-3 days after transfusion, expressed as percentage of KEL+ RBC: WT RBC ratio at the time of transfusion, measured by flow cytometry. Data of one experiment, representative of 3 independent experiments with 4-5 mice per group. *p<0.05 by Kruskal-Wallis test with a Dunn’s post-test.

To determine whether the anti-KEL antibodies were functional, we examined the degree to which anti-KEL antibodies clear transfused RBCs from circulation. Previously transfused mice were re-transfused with fluorescently labeled KEL^+^ and control WT RBCs 35 days after the initial transfusion. Post-transfusion recovery of transfused RBCs was measured by flow cytometry. While approximately half of the KEL+ RBCs transfused to poly(I:C) treated mice were cleared from circulation 3 days after transfusion, post-transfusion recovery of KEL+ RBCs in mice treated with poly(I:C) and CDDO-Im was significantly increased, similar to recovery in WT mice not treated with poly(I:C) (**Figure 1D**). These results indicate that CDDO-Im inhibits production of RBC alloantibodies that mediate clearance of antigen specific RBCs.

### CDDO-Im regulates RBC alloimmunization via Nrf2 activation

Given the possibility that CDDO-Im could alter alloimmunization by Nrf2-independent mechanisms, we examined effects of CDDO-Im on anti-KEL antibody production in Nrf2-deficient (*Nrf2*^-/-^) mice. WT and *Nrf2*^-/-^ were treated with poly(I:C) 3 hrs prior to transfusion. Treatment with CDDO-Im 6 hrs prior to transfusion inhibited anti-KEL IgM and IgG production in WT mice. However, CDDO-Im did not alter anti-KEL IgM and IgG production in *Nrf2*^-/-^ mice (**Figure 2 A,B)**. When examining post-transfusion recovery, *Nrf2*^-/-^ mice treated with and without CDDO-Im had reduced recovery of re-transfused KEL^+^ RBCs, compared to CDDO-Im treated WT mice (**Figure 2C**). These results indicate that CDDO-Im regulates RBC alloimmunization by activating Nrf2.

**Figure 2.**
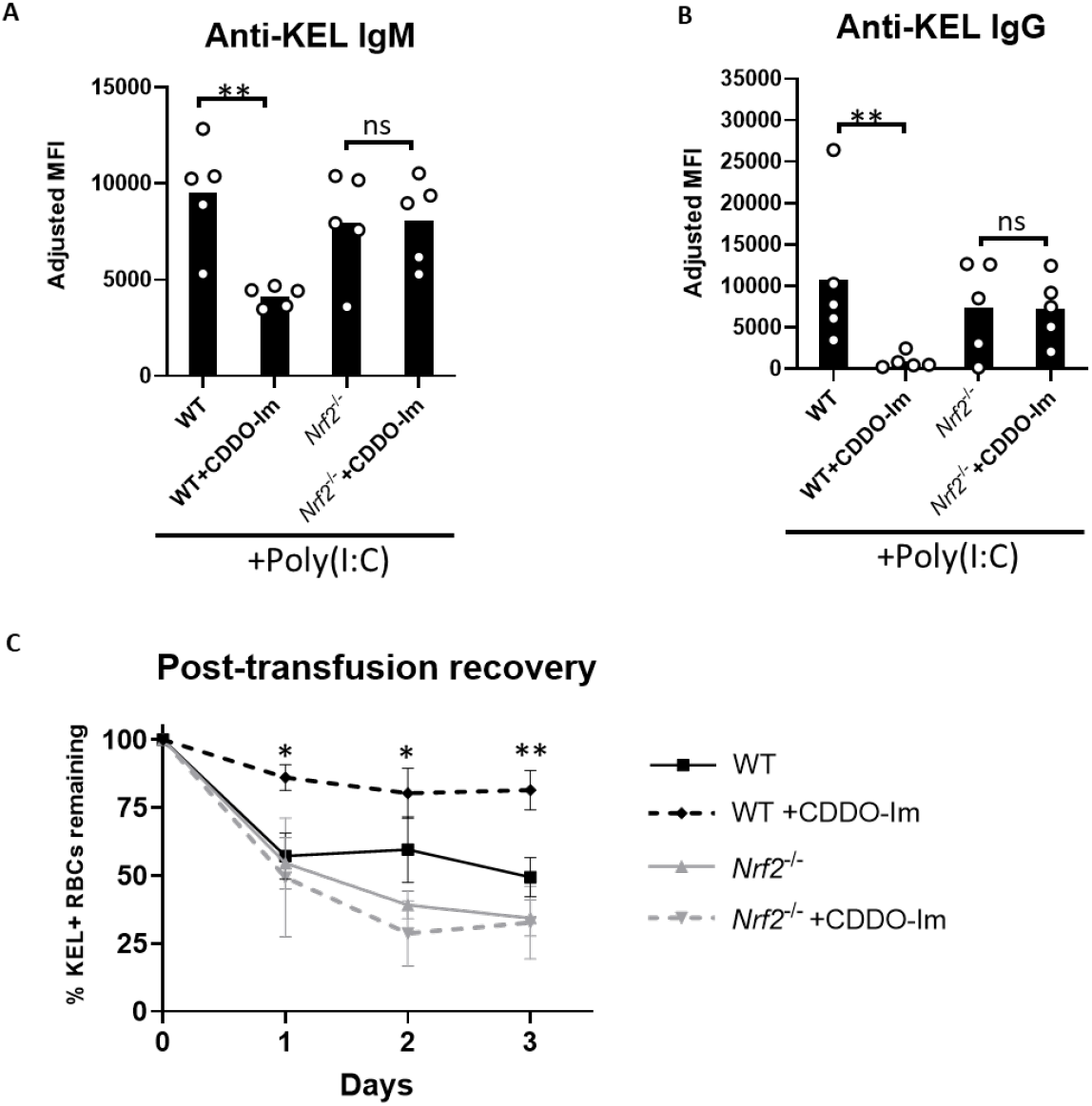
Nrf2 activation inhibits inflammation-induced RBC alloimmunization. WT and *Nrf2*^-/-^ mice were treated with poly(I:C), 3 hrs prior to transfusion with KEL+ RBCs. Mice were treated with or without CDDO-Im 6 hrs before transfusion. (**A**) Serum anti-KEL IgM levels collected 5 days after transfusion. (**B**) Serum anti-KEL IgG levels collected 7 and 14 days after transfusion. (**C**) Mice were re-transfused with fluorescently labeled KEL+ RBCs and control WT RBCs 35 days after the initial transfusion. Post-transfusion recovery of KEL+ RBCs: WT RBCs ratios 1-3 days after transfusion, expressed as percentage of KEL+ RBC: WT RBC ratio at the time of transfusion, measured by flow cytometry. Data of one experiment, representative of 3 independent experiments with 5 mice per group. *p<0.05, **p<0.01 by Kruskal-Wallis test with a Dunn’s post-test.

### CDDO-Im inhibits cytokine production in mice

Nrf2 activation can inhibit production of inflammatory cytokines. Thus, we measured NF-κB-induced cytokines (CXCL1, TNF, IL-6, MCP-1) and IFNα/β in serum of mice treated with CDDO-Im and poly(I:C). CDDO-Im treatment inhibited poly(I:C)-induced CXCL1, TNF, and IL-6, while MCP-1 levels were not significantly affected (**Supplemental Figure 2**). In addition, CDDO-Im inhibited production of poly(I:C)-induced IFNα and IFNβ measured by a multiplex cytokine assay (**Figure 3A**).

**Figure 3.**
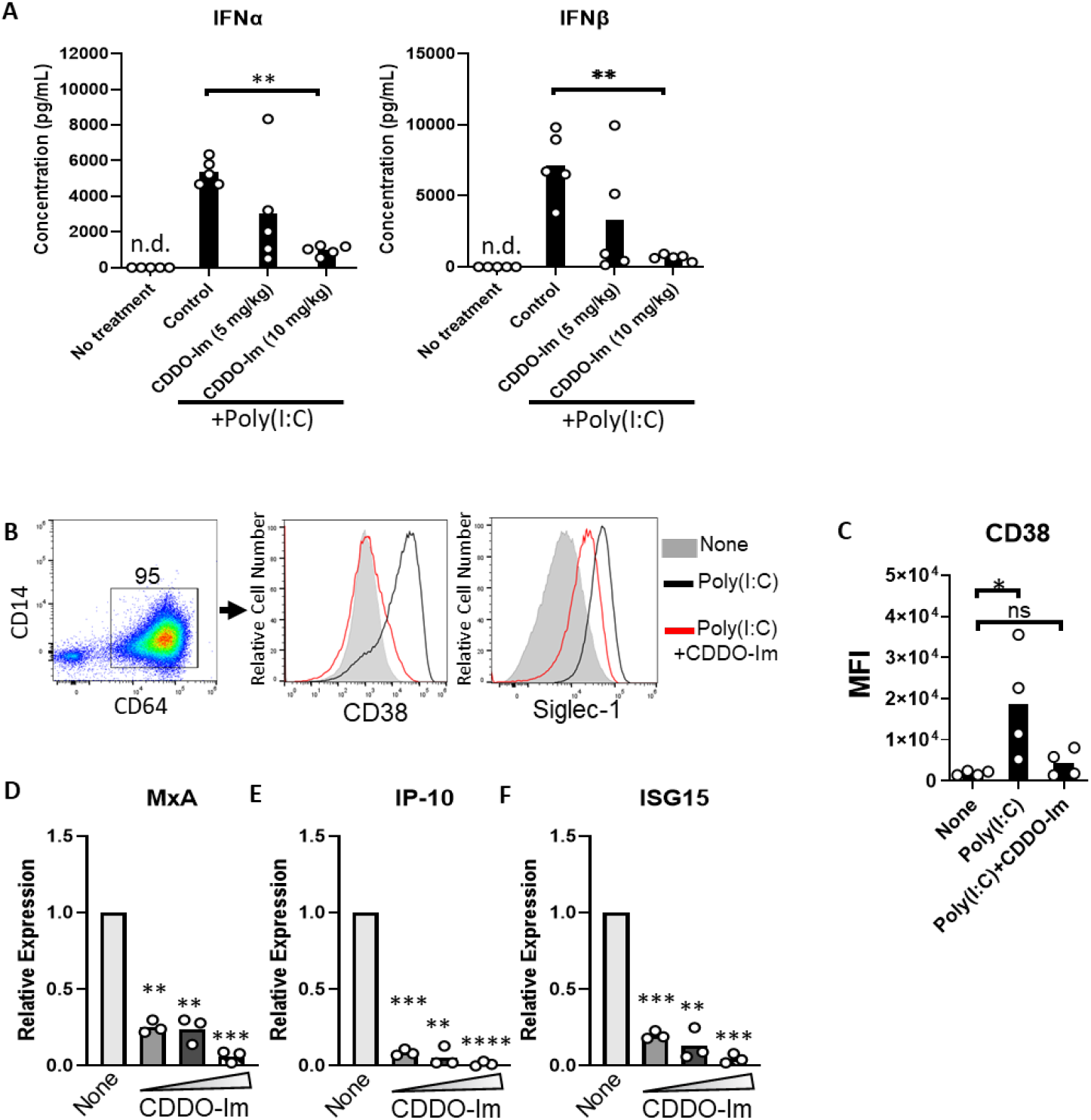
CDDO-Im inhibits IFNα/β activity in mice and human macrophages. **A**) WT mice were treated with poly(I:C) with or without 5-10 mg/kg CDDO-Im 6 hrs and 3 hrs prior to analysis of serum IFNα and IFNβ cytokine levels (HMOX1, NQO1), respectively. **B-F**) Human monocyte-derived macrophages were treated with CDDO-Im for 18 hrs followed by 1 µg/mL poly(I:C) treatment for 3 (**D-F**) or 24 hrs (**B**,**C**). (**B**) Representative flow cytometric analysis of CD38 and Siglec-1 expression on CD64+ macrophages treated with or without poly(I:C) and 0.8 µM CDDO-Im. (**C)** Cumulative data of CD38 expression by macrophages from (**B**). (**D-F**) Expression of ISGs (**D**) MxA, (**E**) IP-10, and (**F**) ISG15 in macrophages treated with poly(I:C) and either 0.2, 0.4, or 0.8 µM CDDO-Im, relative to untreated cells, measured by qPCR. Each circle represents the expression from an independent experiment, n=3. *p<0.05, **p<0.01, ***p<0.001, ****p<0.0001 by one-way ANOVA with a Tukey’s post-test.

### CDDO-Im promotes expression of Nrf2-inducible genes and inhibits IFNα/β signaling in human macrophages

Nrf2 activated genes, including HMOX1 and NQO1, are highly expressed in macrophages, compared to other innate immune cells^35^. Thus, we examined the degree to which CDDO-Im regulates IFNα/β responses in human monocyte-derived macrophages derived from platelet donors. Macrophages were treated with multiple doses of CDDO-Im for 18 hrs. Expression of Nrf2-inducible genes, including AKR1CI, HMOX1, and NQO1 were significantly elevated, compared to untreated macrophages (**Supplemental Figure 3A-C)**. Intracellular flow cytometry showed that Hmox1 protein expression was elevated in CDDO-Im treated CD64^+^ macrophages, compared to controls (**Supplemental Figure 3D, E**).

Given that IFNα/β have been previously shown to induce RBC alloimmunization in murine transfusion models, ISGs were measured in CDDO-Im treated macrophages. Cells were treated with the IFNα/β stimulus, poly(I:C), which increased expression of the ISGs, Siglec-1 and CD38, on the cell surface of CD64^+^ macrophages. However, treatment with CDDO-Im prior to poly(I:C) reduced the expression of Siglec-1 and CD38 (**Figure 3B, C**). In addition, increasing doses of CDDO-Im also inhibited expression of poly(I:C)-induced ISGs, MXA, IP-10, and ISG15, measured by qPCR (**Figure 3D-E**).

To determine whether another Nrf2 activator regulates IFNα/β activity in human macrophages, Nrf2 activated genes and ISGs were measured in macrophages treated with sulforaphane. Like CDDO-Im, sulforaphane increased expression of HMOX1 and NQO1. Following poly(I:C) treatment, prior sulforaphane treatment also inhibited ISG expression (**Supplemental Figure 4**).

### CDDO-Im inhibits RBC alloimmunization in mice with pre-existing inflammation

Given that patients and pre-clinical models with elevated IFNα/β activity have an increased frequency of alloimmunization^9,10,19,36^, we considered whether CDDO-Im also affects alloimmunization in mice with pre-existing IFNα/β activity. Thus, we examined RBC alloimmune responses in WT mice treated with poly(I:C) 3 hrs before CDDO-Im treatment and 6 hrs before transfusion with KEL+ RBCs (**Supplemental Figure 5A)**. At the time of transfusion, IFNα/β and NF-κB-induced cytokine levels were not significantly different between poly(I:C)-treated groups treated with or without CDDO-Im (**Supplemental Figure 5B)**. Following transfusion, flow cytometric crossmatch analysis indicated that CDDO-Im inhibited anti-KEL IgM and IgG production in WT mice previously treated with poly(I:C) (**Supplemental Figure 5C, D**). Re-transfusion with KEL+ and WT RBCs revealed that mice treated only with poly(I:C) prior to the first transfusion had a low post-transfusion recovery of KEL^+^ RBCs. In contrast, mice treated with CDDO-Im after poly(I:C) treatment had a significantly higher recovery of KEL^+^ RBCs (**Supplemental Figure 5E**). Collectively, results indicate that CDDO-Im can suppress alloimmunization in mice with pre-existing inflammation.

## Discussion

Currently, the only strategy to mitigate RBC alloimmunization in transfusion recipients is to match multiple RBC antigens expressed by donors and recipients. However, given that there are hundreds of RBC antigens and many variants of Rh group antigens^7^, alternate approaches are needed to inhibit alloimmunization and subsequent hemolytic events. While significant progress has been made in identifying inflammatory mechanisms that promote RBC alloimmunization^2^, mechanisms that mitigate RBC alloimmunization represent a gap in the field. Here, we report that CDDO-Im-mediated Nrf2 activation inhibits RBC alloimmunization in a pre-clinical transfusion model.

Inflammation induced by the viral mimetic, poly(I:C), induces and enhances the production of alloantibodies against multiple RBC antigens in pre-clinical studies^14^. We have reported that IFNα/β is required for poly(I:C)-induced alloimmunization^17^. In models of lupus and viral infection, mice lacking IFNα/β production or signaling have profoundly reduced alloantibody responses^16,19^. Thus, given that Nrf2 activation was reported to inhibit IFNα/β responses in models of erythrophagocytosis and bacterial infection^23,37^, we tested the degree to which pharmacologic activation of Nrf2 by CDDO-Im regulates IFNα/β responses to poly(I:C). In human macrophages and mouse blood leukocytes, CDDO-Im treatment prior to poly(I:C) treatment inhibited ISG expression and IFNα/β production, respectively. Thus, given the required role for IFNα/β activity in poly(I:C)-induced RBC alloimmunization, CDDO-Im mediated Nrf2 activation may partly regulate alloimmunization by inhibiting IFNα/β activity. However, other contributing factors, including CDDO-Im induced suppression of NF-κB cytokines, cannot be ruled out.

Patients with pre-existing IFNα/β activity, due to autoimmunity, viral infection, or SCD, have an elevated frequency of RBC alloimmunization^9,10,12^. Thus, we tested the degree to which CDDO-Im mediated Nrf2 activity regulates alloimmunization in mice with pre-existing poly(I:C)-induced inflammation. The result that subsequent CDDO-Im treatment inhibited alloimmunization (in the presence of elevated cytokines) indicates that CDDO-Im may also prevent cytokine receptor signaling or have cytokine-independent effects. Nrf2-mediated suppression of IFNα/β-induced STAT1 signaling was previously observed in a model of *Klebsiella pneumoniae* infection, where Nrf2 was induced during erythrophagocytosis of transfused stored RBCs^23^. Further analysis of IFNα/β receptor signaling would determine whether CDDO-Im mediated Nrf2 activation has a dual role of inhibiting IFNα/β production and signaling.

However, it is notable that Nrf2 activation can also regulate other cytokine responses, including those induced by NF-κB (i.e. TNF, IL-6, and CXCL1)^37^. Transfusion of stored RBCs expressing a chimeric RBC antigen has been shown to promote NF-κB-induced cytokines that may enhance RBC alloantibody responses^38^. While this has not been shown for the KEL antigen model reported here, a role for Nrf2 regulation of non-interferon cytokine responses in RBC alloimmunization cannot be ruled out. In addition, it is notable that CDDO-Im alters Nrf2-independent pathways. For example, CDDO-Im regulates apoptosis^39^ and the cell cycle in cancer cells^40^, unfolded protein responses^41^ and other immune pathways including the mammalian target of rapamycin (mTOR) pathway^42^. Thus, it is possible that CDDO-Im off target effects could influence RBC alloimmunization. However, the inability of CDDO-Im to regulate anti-KEL responses in *Nrf2*-deficient mice indicates that CDDO-Im inhibits alloimmunization via Nrf2 activation.

Nrf2-induced genes are highly expressed in spleen macrophages^35^. Specifically, red pulp macrophages continually clear senescent circulating RBCs containing oxidized heme that activates Nrf2^43^. Thus, we examined CDDO-Im effects on IFNα/β activity in human monocyte-derived macrophages. CDDO-Im induced expression of multiple Nrf2 stimulated genes and inhibited poly(I:C)-induced ISG expression, suggesting that immunosuppressive effects of Nrf2 activation can be extended to human macrophages. While macrophages can play a pivotal role in cytokine production and alloimmunization^44^, it is possible that CDDO-Im regulates alloimmunization by regulating other immune cells critical for humoral immune responses. Nrf2 activators have been reported to either promote or inhibit T cell and B cell activation, which may depend on the use of specific activators and disease models^45,46^. Heme-induced Nrf2 activation can also inhibit dendritic cell maturation^43^. Subsequent studies utilizing models of *Nrf2* deficiency or activation in specific cell types will clarify the role of cell-specific Nrf2 activity on RBC alloimmunization.

CDDO-Im is one of many described Nrf2 activating compounds. Dimethy fumurate (DMF) is FDA approved for treatment of multiple sclerosis^29^, and sulforaphane is a Nrf2 activator derived from cruciferous vegetables, including broccoli^28^. While CDDO-Im, DMF, sulforaphane, and many other Nrf2 activators act by altering the principle negative regulator of Nrf2, Keap1, pharmacokinetics, pharmacodynamics, off-target effects, and routes of administration differ. Thus, future studies should delineate the effects of multiple Nrf2 activators on RBC alloimmunization.

Finally, while prophylaxis for RBC alloimmunization may be beneficial in all transfusion recipients, it may be more impactful for patients with SCD, who have the highest frequency of RBC alloimmunization and a high transfusion burden. Interestingly, Nrf2 activators, including CDDO-Im, sulforaphane, and DMF, have been shown to inhibit vaso occlusion and vascular inflammation in models of SCD^32-34^. In addition, a phase 1 trial provided evidence that sulforaphane-containing broccoli sprout homogenates induce expression of multiple Nrf2 target genes in patients with SCD^47^. Thus, prophylaxis for RBC alloimmunization may be one of many beneficial effects of Nrf2 activators in patients with SCD.

In conclusion, we report that the Nrf2 activator, CDDO-Im, inhibits RBC alloimmunization in a Nrf2-dependent manner in a pre-clinical transfusion model. While Nrf2 activation may regulate alloimmunization by multiple mechanisms, CDDO-Im inhibits the IFNα/β response, which promotes inflammation-induced RBC alloimmunization. Findings extending to future studies utilizing alternate Nrf2 activators and other transfusion models may lead to human studies examining the prophylactic potential of currently available Nrf2 activators for transfusion recipients.

## Acknowledgments

Research was supported by the NIH/NHLBI (R01HL165169 to DRG). We thank Stephanie Eisenbarth and Sean Stowell for review and edits of data and figures.

## Authorship and conflict-of-interest statements

Experiments were completed by all authors. Data analysis was performed by C-YC, RHA, KP, KN, and DRG. DRG drafted the manuscript, which was reviewed and edited by all authors. The authors have no competing financial interests.

