## Supplemental Material for "CDDO-Imidazole regulates RBC alloimmunization to the KEL antigen by activating Nrf2"

**Supplemental Table 1. Primer Sequences used for quantitative PCR.**

| Primer Name | Forward | Reverse |
|---|---|---|
| Mouse HMOX1 | 5'- AGG CTA AGA CCG CCT TCC T -3' | 5'- TGT GTT CCT CTG TCA GCA TCA -3' |
| Mouse *NQO1* | 5'- GCC GAA CAC AAG AAG CTG GAA G -3' | 5'- GGC AAA TCC TGC TAC GAG CAC T -3' |
| Mouse GAPDH | 5'- CAT CAC TGC CAC CCA GAA GAC TG -3' | 5'- ATG CCA GTG AGC TTC CCG TTC AG -3' |
| Human *NFE2L2* (Nrf2) | 5'- CAG CGA CGG AAA GAG TAT GA -3' | 5'- TGG GCA ACC TGG GAG TAG -3' |
| Human AKR1C1 | 5'- CGA GAA GAA CCA TGG GTG GA -3' | 5-' GGC CAC AAA GGA CTG GGT CC -3' |
| Human *HMOX1* | 5 -GCT GCT GAC CCA TGA CAC CAA GG-3 | 5 -AAG GAC CCA TCG GAG AAG CGG AG-3 |
| Human *NQO1* | 5'- GAA GAG CAC TGA TCG TAC TGG C-3' | 5'- GGA TAC TGA AAG TTC GCA GGG -3' |
| Human *MXA* | 5'- CTC CGA CAC GAG TTC CAC AA -3' | 5'- GGC TCT TCC AGT GCC TTG AT -3' |
| Human *CXCL10* (IP-10) | 5-' GGT GAG AAG AGA TGT CTG AAT CC -3' | 5-' GTC CAT CCT TGG AAG CAC TGC A -3' |
| Human ISG15 | 5'- ACT CAT CTT TGC CAG TAC AGG AG -3' | 5'- CAG CAT CTT CAC CGT CAG GTC -3' |
| Human IFIT3 | 5'- AGA GAC ACA GAG GGC AGT CA -3' | 5-' GGC ATT TCA GCT GTG GAA GG -3' |
| Human IRF5 | 5'- TAT GCC ATC CGC CTG TGT CAG T -3' | 5'- GCC CTT TTG GAA CAG GAT GAG C- 3' |
| Human IRF7 | 5'- AAA ACC AAC TTC CGC TGC -3' | 5'- GCC TCA GTC TGG TCC GTG C-3' |
| Human GAPDH | 5-' TCA CCA GGG CTG CTT TTA AC -3' | 5'- ACA AGC TTC CCG TTC TCA G-3' |

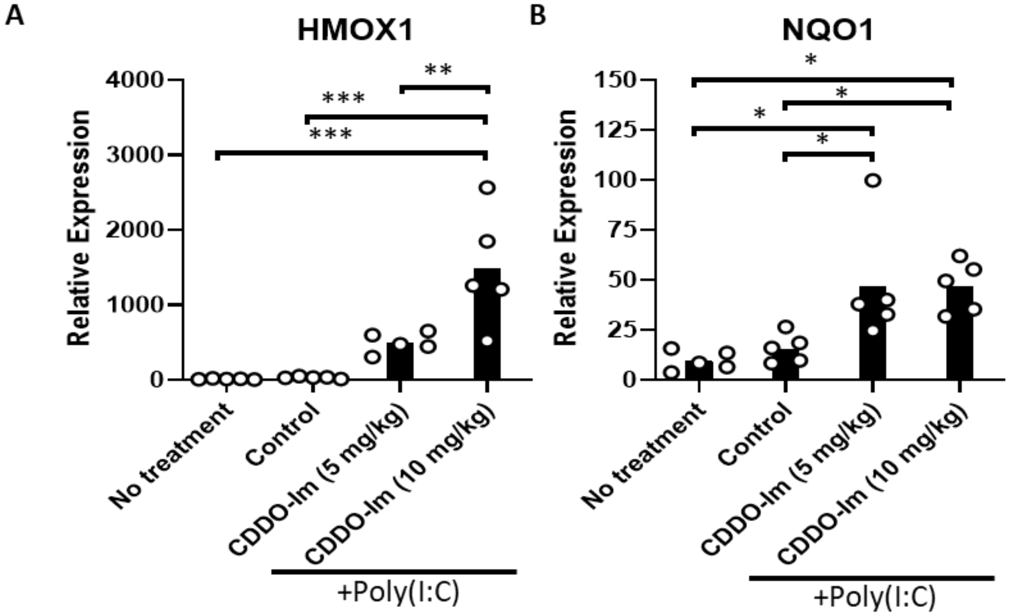


**Supplemental Figure 1. CDDO-Im induces Nrf2-activated gene expression in mice.** WT mice were treated (i.p.) with 5-10 mg/kg CDDO-Im and/or 100 µg poly(I:C) 6 hrs and 3 hrs prior to analysis, respectively. (**A**) HMOX1 and (**B**) NQO1 expression by blood leukocytes measured by quantitative PCR. *p<0.05, **p<0.01, ***p<0.001 by One-way ANOVA with a Tukey's post-test; 5 mice per group; representative of 2 independent experiments.

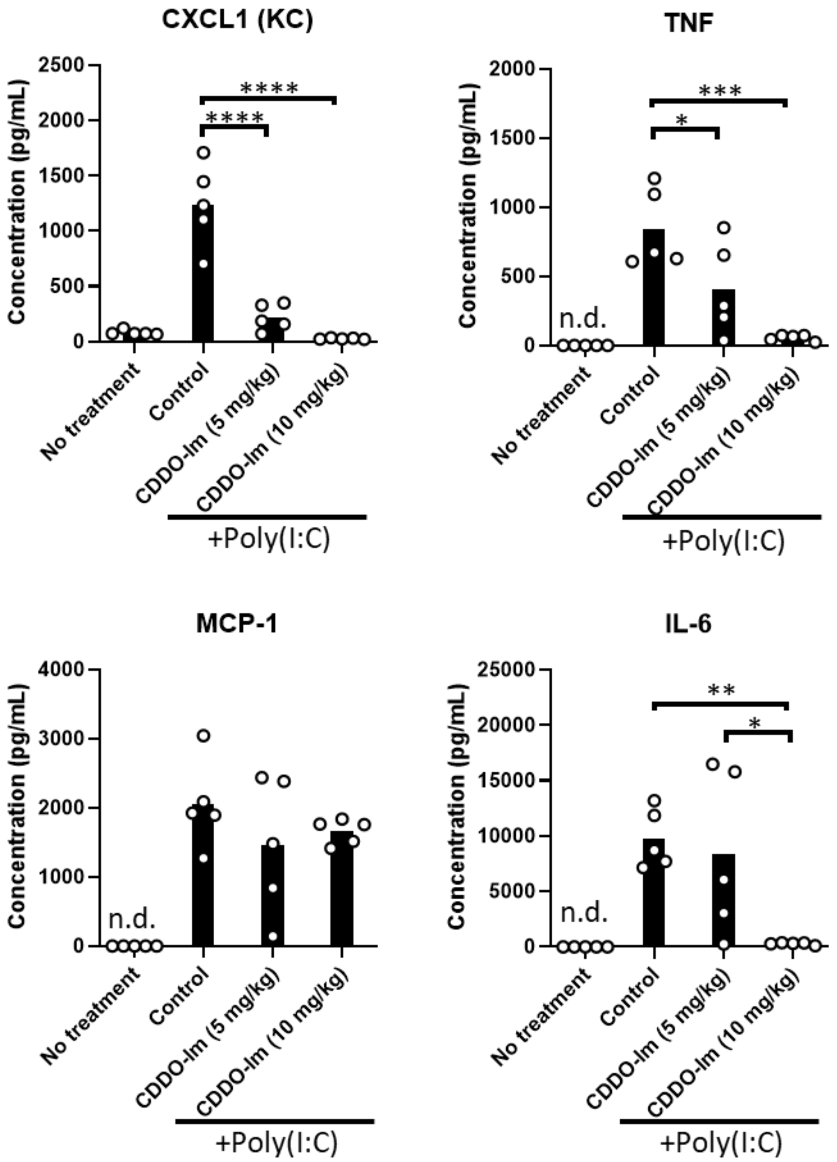


**Supplemental Figure 2. CDDO-Im regulates NF-κB-induced cytokines in mice.** WT mice were treated (i.p.) with 5-10 mg/kg CDDO-Im and/or 100 µg poly(I:C) 6 hrs and 3 hrs prior to analysis, respectively. CXCL1, TNF, MCP-1, and IL-6 levels in serum measured by multiplex bead assay. *p<0.05, **p<0.01, ***p<0.001, p<0.0001 by One-way ANOVA with a Tukey's post-test; 5 mice per group; representative of 2 independent experiments; n.d. = not detectable.

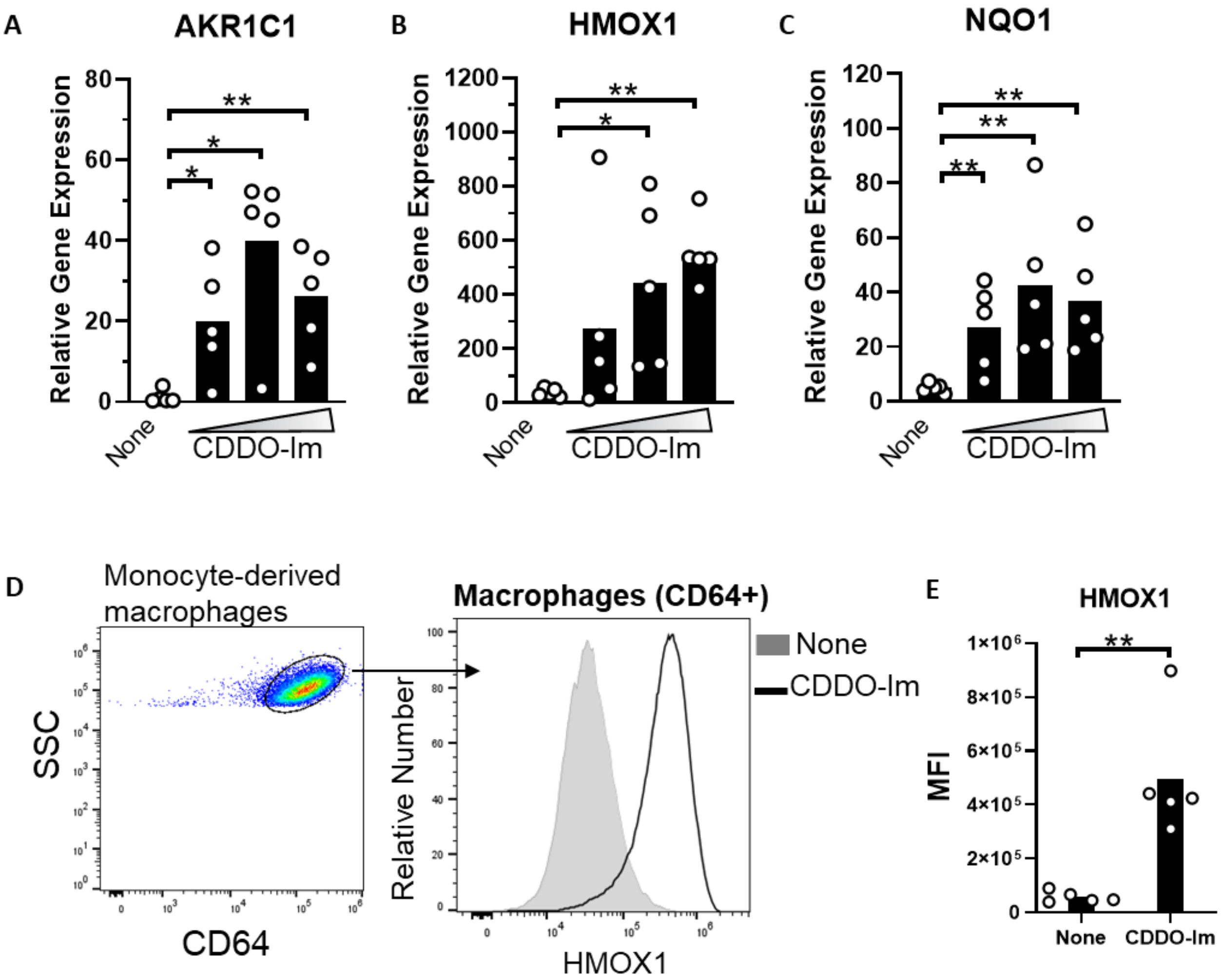


**Supplemental Figure 3. CDDO-Im induces Nrf2-activated gene expression in human macrophages.** Human monocyte-derived macrophages were treated with CDDO-Im for 18 hrs. (**A**) AKR1C1, (**B**) HMOX1, and (**C**) NQO1 expression in macrophages treated with either 0, 0.2, 0.4, or 0.8 µM CDDO-Im, measured by qPCR (**D**) Representative intracellular flow cytometric analysis of HMOX1 expression by CD64+ macrophages treated with or without 0.8 µM CDDO-Im. **(E)** Cumulative data of HMOX1 expression by macrophages from 5 independent experiments; **p<0.01 by Student's t-test. (**D**). (**A-C**) Each circle represents the expression from an independent experiment, n=5. *p<0.05, **p<0.01, ***p<0.001, ****p<0.0001 by one-way ANOVA with a Tukey's post-test. (**D**) Representative of 5 independent experiments.

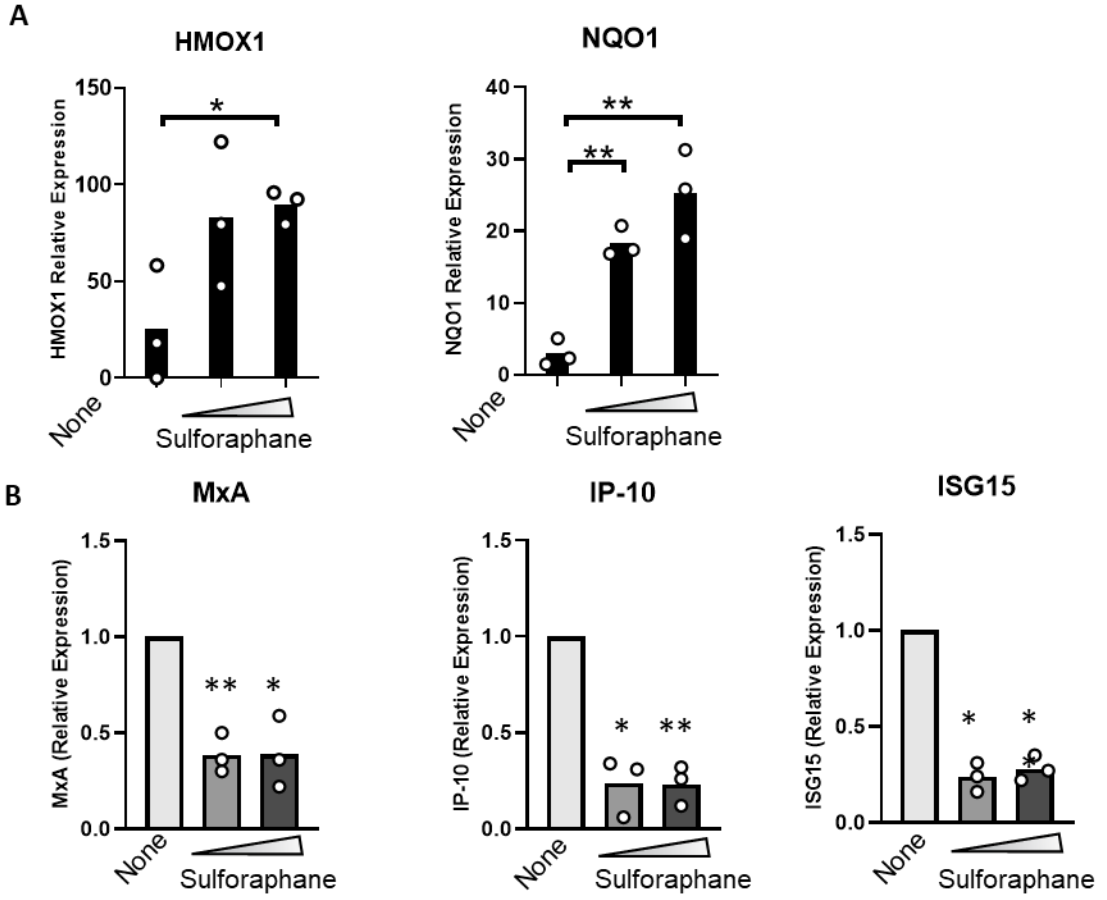


**Supplemental Figure 4. Sulforaphane induces Nrf2-activated gene expression and regulates IFNα/β activity in human macrophages.** Human monocyte-derived macrophages were treated with 0, 5, or 10 μM for 18 hrs. (**A**) Expression of Nrf2-activated genes, HMOX1 and NQO1, by qPCR. (**B**) Following sulforaphane treatment, macrophages were treated with poly(I:C) for 3 hrs. Macrophage expression of ISGs, MxA, IP-10, and ISG15, relative to macrophages not treated with CDDO-Im, measured by qPCR. *$p<0.05$, **$p<0.01$ by one-way ANOVA with a Tukey's post-test. Each circle represents an independent experiment, n=3.

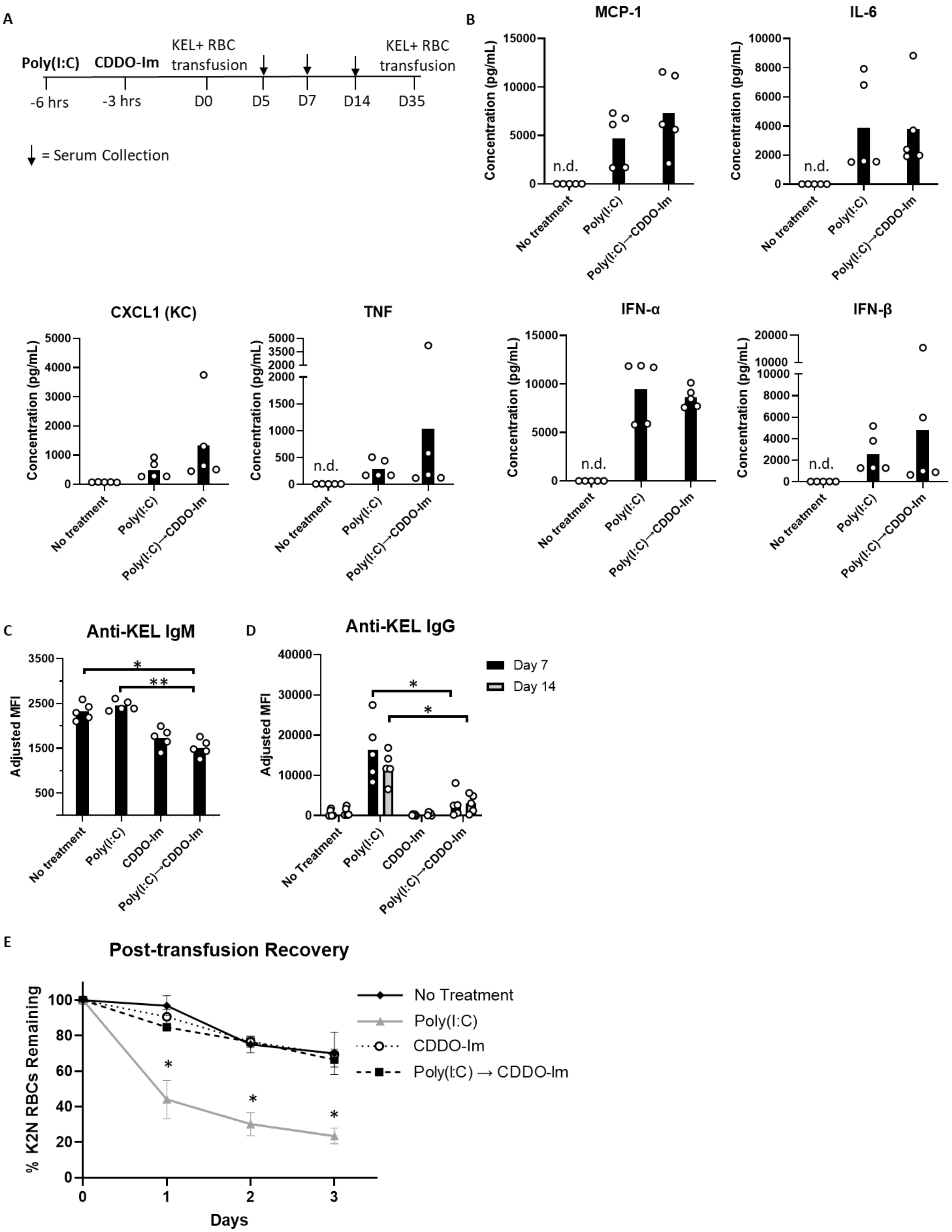

A
Poly(I:C)
CDDO-Im
KEL+ RBC transfusion
KEL+ RBC transfusion
-6 hrs
-3 hrs
D0
D5
D7
D14
D35
= Serum Collection
B
MCP-1
IL-6
CXCL1 (KC)
TNF
IFN-α
IFN-β
Concentration (pg/mL)
n.d.
No treatment
Poly(I:C)
Poly(I:C)→CDDO-Im
C
Anti-KEL IgM
Adjusted MFI
No treatment
Poly(I:C)
CDDO-Im
Poly(I:C)→CDDO-Im
D
Anti-KEL IgG
Day 7
Day 14
Adjusted MFI
No Treatment
Poly(I:C)
CDDO-Im
Poly(I:C)→CDDO-Im
E
Post-transfusion Recovery
% K2N RBCs Remaining
Days
No Treatment
Poly(I:C)
CDDO-Im
Poly(I:C) → CDDO-Im

**Supplemental Figure 5. CDDO-Im regulates RBC alloimmunization in mice with pre-existing inflammation.** WT mice were treated with or without poly(I:C) and/or CDDO-Im 6 hrs and 3 hrs before transfusion, respectively, with KEL+ RBCs. (**A**) Timeline of treatments, transfusions, and serum collections. (**B**) Serum cytokines at the time of transfusion. (**C**) Serum anti-KEL IgM levels measured 5 days after transfusion. (**D**) Serum anti-KEL IgG levels measured 7 and 14 days after transfusion. (**E**) Mice were re-transfused with fluorescently labeled KEL+ RBCs and control WT RBCs 35 days after the initial transfusion. Post-transfusion recovery of KEL+ RBCs: WT RBCs ratios 1-3 days after transfusion, expressed as percentage of KEL+ RBC: WT RBC ratio at the time of transfusion, measured by flow cytometry. Data of one experiment representative of 2 independent experiments with 5 mice per group. $^{*}p<0.05$, $^{**}p<0.01$ by Kruskal-Wallis test with a Dunn's post-test.
